# MyD88-dependent signaling promotes smooth muscle cell phenotypic modulation and fibrous cap formation in murine atherosclerosis

**DOI:** 10.1101/2025.07.01.662493

**Authors:** Stine Gunnersen, Julián Albarrán-Juárez, Lise Filt Jensen, Jeong Tangkjær Shim, Charlotte B. Sørensen, Jacob F. Bentzon

## Abstract

**Background and aims:** Recruitment of fibrous cap smooth muscle cells (SMCs) is critical for stabilizing atherosclerotic plaques and preventing rupture. This study investigated the SMC-specific role of the myeloid differentiation primary response protein 88 (*Myd88*) gene, encoding an adaptor protein essential for signaling downstream of several cytokine and pattern-recognition receptors, in this process.

**Methods:** The effects of *MyD88* knockdown were assessed in cultured rat aortic SMCs. SMC-specific knockout of *Myd88* was induced in mice using the Cre/lox method, followed by induction of atherosclerosis by proprotein convertase subtilisin/kexin type 9 gene transfer and a high-fat diet for 12 or 20 weeks. Effects on plaque and fibrous cap formation were studied by immunofluorescence and in situ hybridization.

**Results:** *Myd88* knockdown reduced proliferation and migration of cultured SMCs and preserved contractile gene expression under inflammatory stimulation. SMC-specific *Myd88* deficiency in hyperlipidemic mice did not significantly alter plaque size in the aortic root but reduced the number of cap SMCs in advanced lesions at 20 weeks and in the most atherosclerosis-susceptible aortic sinus at 12 weeks. Other plaque features, including macrophages, necrotic core size, and collagen content, were not significantly affected. Notably, MyD88 deficiency preserved the contractile phenotype of medial SMCs beneath plaques, suggesting that impaired phenotypic modulation contributed to reduced cap SMC recruitment.

**Conclusions:** MyD88-dependent signaling promotes medial SMC phenotypic modulation and the recruitment of fibrous cap SMCs during atherogenesis. These findings highlight MyD88 as a mediator linking inflammatory signaling to protective fibrous cap formation.

**Highlights:**

- Knockdown of *MyD88* preserves the contractile phenotype of cultured smooth muscle cells subjected to inflammatory cytokines.
- Targeting *MyD88* in murine atherosclerosis inhibits fibrous cap formation.
- Targeting *MyD88* in murine atherosclerosis preserves the contractile phenotype of medial smooth muscle cells beneath the plaque.

## Introduction

The fibrous cap of atherosclerotic lesions is produced by smooth muscle cells (SMCs) that assemble underneath the endothelium and generate a fibrous extracellular matrix.^1^ The cap recreates mechanical support for the endothelium, which is temporarily lost in early atherosclerosis by the interposition of macrophages and other cell types between the endothelium and the arterial media. Its thickness and strength are critical for protecting plaques against rupture, which is the most common cause of thrombosis, leading to life-threatening heart or brain infarction.^2^ Understanding the mechanisms involved in fibrous cap formation is therefore important for developing therapies to mitigate the harmful consequences of atherosclerotic plaque formation.

In mouse models of atherosclerosis, most cap SMCs derive from a few clonally expanding medial SMCs, which form a layered structure in the subendothelial space orchestrated by Notch and PDFG signaling.^3–6^ It remains unknown, however, how the medial SMC founders are recruited to this process. One hypothesis consistent with existing knowledge is that inflammatory cytokines and damage-associated molecular patterns generated in early atherosclerosis may activate the founder SMCs through NFκB signaling. Canonical NFκB signaling leads to SMC de-differentiation and proliferation in vitro,^7^ and deleting the essential NFκB signaling protein IKKβ in SMCs inhibits atherogenesis in mice.^8^ Furthermore, we recently found that atherosclerotic disease activity is tightly associated with NFκB-controlled gene expression in SMC-derived plaque cells.^9^

There are several parallel pathways through which NFκB signaling in SMCs can be activated in early atherosclerosis. Toll-like receptors (TLRs) constitute a large family of NFκB-activating pattern-recognition receptors that bind pathogen- and damage-associated molecular patterns, including those formed on modified ApoB-LPs.^10^ Global knockout of *Tlr2* and *Tlr4* inhibits plaque development in *Apoe-/-* mice,^11,12^ and the fact that *Tlr2* deletion in hematopoietic cells did not have the same effect, suggests that it is partly mediated by SMCs. The interleukin-1 receptor type 1 (IL1R1) that binds the pro-inflammatory cytokine IL-1β also signals through the NFκB signaling pathway and is important for both murine and human atherosclerosis.^13–15^ Importantly, much of the effect on plaque development in mice appears to be mediated by IL1R1s in SMCs based on the analysis of cell-type specific *Il1r1* knockouts.^15,16^

Considering the range of receptors and ligands that impinge on NFκB signaling, significant redundancy is expected. However, broad inhibition of signaling from most TLRs and the IL1R1 can be achieved by targeting the adaptor protein MyD88 (Myeloid differentiation primary response protein 88).^17^ Upon IL-1R/TLR stimulation, MyD88 is recruited to the IL1R1/TLR through binding of the TIR domain, which mediates the binding of IL-1 receptor-associated kinase (IRAK) with the DD domain.^18^ This triggers IRAK phosphorylation, which is crucial for inducing a signaling cascade of events leading to NFκB activation.^18,19^ Global MyD88 deficiency reduced murine atherosclerosis,^11,20^ but the isolated effect in SMCs is not known. In the present study, we analyzed the role of MyD88-dependent signaling in SMCs for atherogenesis and cap SMC recruitment in hypercholesterolemic mice. We find that blocking MyD88-dependent signaling mitigates de-differentiation of medial SMCs and inhibits the accumulation of cap SMCs in plaques.

## Materials and methods

### Cell culture

Rat aortic smooth muscle cells (#CRL-2018, ATCC) were cultured in DMEM medium supplemented with 10% FBS, 1% penicillin/streptomycin, and 2 mM glutamine, in a humidified incubator at 37 °C with 5% CO2. For all experiments, cells were seeded in 24-well plates coated with collagen type I (5 mg/ml). At 70% confluence, the cells were transfected twice with 10 µM control siRNA (AllStars Negative siRNA, #1027280, Qiagen) or 10 µM of *Myd88* siRNAs (#SI03063431 and #SI03076885, Qiagen) using Opti-MEM™ I (#31985062, Thermo Fisher Scientific) and Lipofectamine RNAiMAX transfection reagent (#13778030, Thermo Fischer Scientific) for 18 h on two consecutive days. Cells were then treated with 10 ng/ml recombinant human IL1B protein (#PHC0814, Thermo Fisher Scientific), 100 ng/ml LPS (#tlrl-eklps, InvivoGen), or vehicle for 24h. Each experiment contained 4 replicates per condition.

To analyze gene expression changes, total RNA was isolated using the Nucleospin RNA Plus mini kit according to the manufacturer’s protocol (#740984.50, Macherey Nagel). Complementary DNA synthesis was performed using 0.5 μg of RNA with the RevertAid First Strand cDNA synthesis kit (#K1622, Thermo Fischer Scientific). For Quantitative real-time PCR, 100 ng of cDNA served as the template for PCR amplification with primers in **Supplemental Table 1** using the Maxima SYBR Green/ROX QPCR Master Mix (2x) according to the manufacturer’s instructions (#K0221, Thermo Fischer Scientific) running on an Aria Mx3000P qPCR System (Agilent Technologies, Santa Clara, CA). Each reaction was run in duplicates, and relative gene expression levels were averaged and normalized to rat hypoxanthine-guanine phosphoribosyltransferase (*Hprt1*). Relative expression was calculated using the ΔΔCt method.

To analyze migration, a siRNA-transfected confluent cell monolayer was scratched in the center of the well using a 10 μl pipette tip before washing with PBS to clear cell debris and floating cells. The cells were incubated for 24 h at 37 °C in 5% CO_2_ using an Eclipse Ti2 inverted microscope (Nikon) with an integrated chamber. Time-lapse images were captured under the microscope every hour for 72 h at the same position. Migration ability was measured by calculating the percentage of area covered by cells after 8 h compared with the area at time 0 using NIS-Elements software (Nikon).

To analyze proliferation, siRNA-transfected cells were cultured for 24 h in DMEM medium containing 10 µM EdU (BCK-EDU488, Sigma-Aldrich). Cells were fixed in 3.7% paraformaldehyde in phosphate-buffered saline (pH 7.4; PBS) for 15 min. The EdU-positive cells were labeled according to the manufacturer’s instructions, followed by immunofluorescence labeling of nuclei with DAPI. Stained coverslips were mounted on slides and analyzed using fluorescence microscopy with a 10X objective. Nikon NIS-Elements software was used for image acquisition, and ImageJ (NIH) was used for quantification.

### Mice

All animal procedures were approved by The Danish Animal Experiments Inspectorate (2020-15-0201-0411) and conducted at Aarhus University. Male *Myh11-*CreER^T2^ mice (B6.FVB-Tg(Myh11-cre/ERT2)1Soff/J, The Jackson Laboratory stock #019079), expressing tamoxifen-inducible *CRE* under SMC-specific *Myh11* promoter,^21^ were crossed with female *Myd88*^*flx*^ mice (B6.129P2(SJL)-Myd88^tm1Defr^/J, The Jackson Laboratory stock #008888) with *loxP* sites flanking exon 3 of the *Myd88* gene.^22^ Male *Myh11*-CreER^T2^ x *Myd88*^flx/flx^ mice (*Myd88*^SMC-KO^) and *Myh11*-CreER^T2^ x *MyD88*^wt/wt^ (*Myd88*^SMC-WT^) littermates were injected with tamoxifen (#T5648, Sigma-Aldrich) dissolved in peanut oil (#P2144, Sigma-Aldrich), 1 mg i.p. daily for 2 × 5 days separated by 2 days without injections, at 6-8 weeks of age. Four weeks after the last tamoxifen injection, atherogenic hypercholesterolemia was induced by a single tail vein injection of rAAV8-mD377Y-mPCSK9 virus particles (1 × 10^11^ vector genomes, produced at Vector Core at the University of North Carolina, Chapel Hill, North Carolina, USA) combined with feeding a high-fat diet (#D12079B, Research Diets Inc.) containing 21% fat and 0.2% cholesterol.^23^ Blood samples were collected from the submandibular vein 2, 6, 11, and 19 weeks after rAAV8-PCSK9 injection. Non-fasting total plasma cholesterol was measured in duplicates with an enzymatic reagent (#CH201, Randox). Mice were anesthetized with 5 mg pentobarbital with 10% lidocaine, killed by exsanguination at 12 (n=22 *Myd88*^SMC-KO^ and n=19 *Myd88*^SMC-WT^) or 20 weeks (n=9 *Myd88*^SMC-KO^ and n=14 *Myd88*^SMC-WT^) after rAAV8-PCSK9 injection, and perfused through the left ventricle with Cardioplex (30 seconds) and 4% phosphate-buffered formaldehyde (5 minutes) at 100 mmHg. Mice were immersed in 4% phosphate buffered formaldehyde for 24 hours and then stored in phosphate-buffered saline at 4°C until tissue harvesting. A control set of *MyD88*^flx/flx^ mice (n=15, without Cre recombinase) were treated similarly with 12 weeks of atherogenesis to allow the calculation of *MyD88* recombination efficiency.

### Aortic root sectioning and staining

The upper part of the heart was dehydrated and embedded in paraffin. Aortic root cross-sections were collected starting at the level of the commissures of the aortic valve and serially sampled at 80 µm intervals from this level. Unfolded sections in the correct orientation were not obtained from 7 mice in the 12-week and 2 mice in the 20-week experiment due to technical problems. Sections from levels 0 µm, 80 µm, 160 µm, and 240 µm were stained by hematoxylin-eosin, and the atherosclerotic plaque area was quantified in micrographs using ImageJ (Fiji distribution).^24^ Separate sections at the 0 µm level were stained with Picrosirius Red and analyzed with polarization microscopy to visualize collagen and to aid measurement of the necrotic core area, which was defined as the absence of collagen and cells (the latter judged from adjacent hematoxylin-eosin-stained sections). Immunofluorescence was performed on sections close to the aortic valve commissures with 1) rat monoclonal IgG2a anti-Mac2 (#CL8942AP, Cederlane, 1:200) followed by Alexa 488-conjugated donkey anti-rat (#A21208, Invitrogen, 1:400); 2) rabbit anti-CD68 (#ab125312, Abcam, 1:200) followed by Alexa 647 donkey anti-rabbit (#ab150075, Abcam, 1:400), 3) mouse monoclonal anti-ACTA2 (#M0851, Dako, 1:100) after blocking endogenous mouse IgG with anti-mouse Fab fragments (#715-007-003, Jackson ImmunoResearch) and followed by Alexa 488-conjugated donkey anti-mouse (#A21202, Invitrogen, 1:400) secondary antibodies, and 4) multiple anti-MyD88 antibodies as listed in **Supplemental Table 2**. DAPI was added to secondary antibodies to visualize nuclei, coverslips were mounted with Slowfade™ Gold Antifade (#S36936, Invitrogen), and images were acquired on an Olympus CellR fluorescence microscope and analyzed in ImageJ (Fiji distribution) using the ROI tool (plaque size, necrotic core size), threshold tool (CD68), and by manual counting of positive nucleated cells (ACTA2, LGALS3).

Staining specificity was validated by substituting the primary antibody with an isotype control. For anti-MyD88 antibodies, specificity was further tested on tissues from global *Myd88* knockout (generated by germline recombination of the floxed Myd88 strain, stock #009088) and control C57BL/6J mice (stock #000664) from The Jackson Laboratory.

### RNAscope in situ hybridization

*MyD88* and *Acta2* RNA detection was performed on aortic root sections (at the 80 μm level) using RNAscope^®^ (RNAscope^®^ Multiplex Fluorescent V2 Reagent kit, #323100, Biotechne) per the manufacturer’s protocol with a mouse *MyD88* probe (#528891, Biotechne), recognizing residues between the region 333-1260 bp of the *Myd88* transcript (accession number NM_010851.2), and a mouse *Acta2* probe (#319531-C2, Biotechne), with a target region between 41-1749 bp of the *Acta2* transcript (NM_007392.3). Both RNAs were visualized with Opal™ 570 (#FP1488001KT, Akoya Biosciences) and nuclei were labeled with DAPI. Slides were mounted in ProLong™ Gold Antifade Mountant (#P10144, Thermo Fischer Scientific). Individual images were collected using a Nikon Eclipse Ti2 fluorescent microscope. For all analyses, images were captured with a 40X Plan Apo l (NA 0.9, WB 250m), which enabled the identification of foci corresponding to individual transcripts. 3 to 5 images were collected per vessel cross-section from 5-6 mice per genotype. The arterial media in each sample was defined and selected in ImageJ (Fiji distribution). For RNA quantification on each sample, the fluorescence intensity was measured.

### En face quantification

For *en face* quantification of atherosclerosis, the aortic arch (down to the supreme intercostal artery) was dissected from the mouse. All periadventitial fat was removed and the aorta was cut open longitudinally, stained with Oil Red O (#O0625, Merck, 3 mg/ml) for 10 minutes at 37^°^C and mounted on a microscope slide with AquaTex (#1.08562, Merck). Three aortas from the 12-week experiment had to be excluded for technical reasons. Aortic arch lesion coverage was quantified by measuring plaque area as a percentage of total aorta area. The slides were scanned with the Epson Perfection V600 Photo scanner.

### Recombination efficiency

Recombination efficiency was determined in normal segments of the aorta from *Myd88*^SMC-KO^ and Cre-negative *Myd88*^flx/flx^ mice after 12 weeks of atherogenesis. The adventitia was removed, the endothelium was scraped off with a scalpel, and DNA was extracted from the remaining arterial medial layer using QIAamp DNA FFPE Tissue Kit (#56404, Qiagen). Quantitative PCR was performed with Maxima SYBR Green/ROX qPCR Mastermix (2x) (#K0221, Thermo Fischer Scientific) and *Myd88*-specific primers (5’-CGATGCCTTTATCTGCTA-3’, 5’-CATTACCCAGGCGGGACAG-3’) flanking the downstream loxP site of the non-recombined floxed Myd88 allele. The *Actb* gene was used for normalization using the primers 5’-CAGCCAACTTTACGCCTAGC-3’ and 5’-TCTCAAGATGGACCTAATACGG-3’. Cycling conditions were: 1 cycle of 10 minutes at 95^°^C followed by 40 cycles of 30 seconds at 95^°^C, 2 minutes at 66^°^C (for *Myd88)* or 66^°^C (for *Actb*), 1 minute at 72^°^C and finally, 1 cycle of 1 minute at 95^°^C, 30 seconds at 60^°^C and 30 seconds at 95^°^C. The ΔΔCt method was used to calculate fold change and recombination efficiency.

### Statistics

Statistical analysis was performed using Prism (GraphPad Software Inc.). Normal distribution was assessed using D’Agostino-Pearson omnibus normality test. Two-group comparisons were performed using unpaired t-tests, Mann-Whitney, or repeated-measures ANOVA as indicated. *P* <0.05 was considered statistically significant. Error bars represent standard errors of the mean (SEM) as indicated in each figure legend.

## Results

### Knockdown of *Myd88* in cultured SMCs reduces phenotypic modulation

To analyze the importance of MyD88-dependent signaling in SMCs, we knocked down *Myd88* in cultured rat aortic SMCs using siRNAs and analyzed responses to 24 hours of interleukin-1β (IL1B) and lipopolysaccharide (LPS) exposure, which activate NFκB signaling through the IL1R1 and TLR4, respectively. *Myd88* expression was reduced by more than 90% compared with cells treated with control siRNA (**Fig. 1A**), and the loss blocked IL1B and LPS-induced increases in inflammatory gene expression (*Vcam1* and *Il6*) (**Fig. 1B-C**). Interestingly, both proinflammatory stimulants downregulated markers of the contractile SMC phenotype (*Myh11, Acta2*, and *Smnt*), and this effect required MyD88 as it was almost abolished by the *MyD88* knockdown (**Fig. 1D-F**). Furthermore, we found that loss of Myd88 significantly lowered migration and proliferation, an effect present both with and without IL1B and LPS stimulation (**Fig. 1G-H**). Overall, the findings were consistent with the potential role of MyD88-dependent signaling in linking the pro-inflammatory cytokines and damage-associated molecular patterns in atherosclerosis to the de-differentiation, migration, and proliferation of local SMCs.

**Figure 1.**
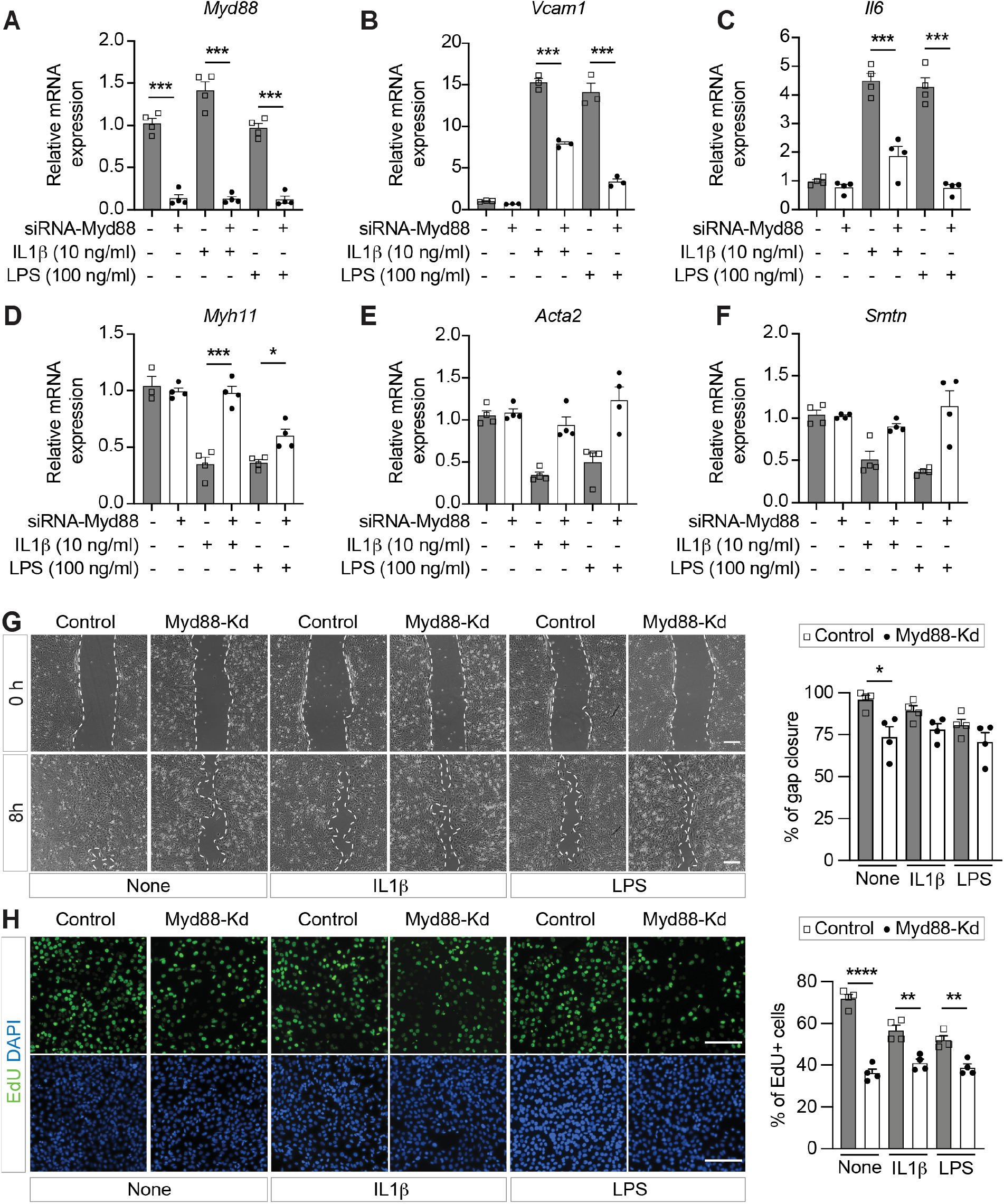
*Myd88* knockdown in cultured SMCs inhibits phenotypic modulation. **A-F**. Gene expression analysis of proinflammatory (A-C: *Myd88, Vcam1, Il6)*, and contractile (D-F: *Myh11, Acta2*, and *Smtn)* gene markers in rat SMCs. Cells were transfected with control or Myd88 siRNAs and then left untreated (none), stimulated with IL1β (10 ng/ml) or LPS (100 ng/ml) for 24 h. Four replicates were evaluated for each condition. Statistical analysis was performed with ANOVA followed by Dunnett′s test; *P <0.05, ***P <0.001. **G-H**. Scratch assay (G) and cell proliferation analysis (H) were performed in confluent rat SMCs monolayers. The percentage of gap closure (G, after 8h) or EdU-positive cells (H, after 24h) was calculated in control and Myd88-Kd SMCs. Nuclei were labeled with DAPI. Four replicates were evaluated for each condition. Scale bars in G-H, 200 μm. Groups were compared by unpaired t-tests; *P <0.05, ***P <0.001.

### Mice with SMC-specific knockout of *Myd88*

To analyze the role of SMC-expressed MyD88 for atherosclerosis development, we generated littermate mice with SMC-restricted CreER^T2^ expression and either homozygous floxed (*Myd88*^SMC-KO^) or wildtype *Myd88* alleles (*Myd88*^SMC-WT^) (**Fig. 2A**). Recombination was induced by tamoxifen at 6 weeks of age. Atherogenesis was initiated at 12 weeks of age via intravenous injection of rAAV8-PCSK9 followed by feeding on a high-fat diet (**Fig. 2B**). Two experiments were performed with atherosclerotic lesions examined after 12 or 20 weeks of disease progression.

**Figure 2.**
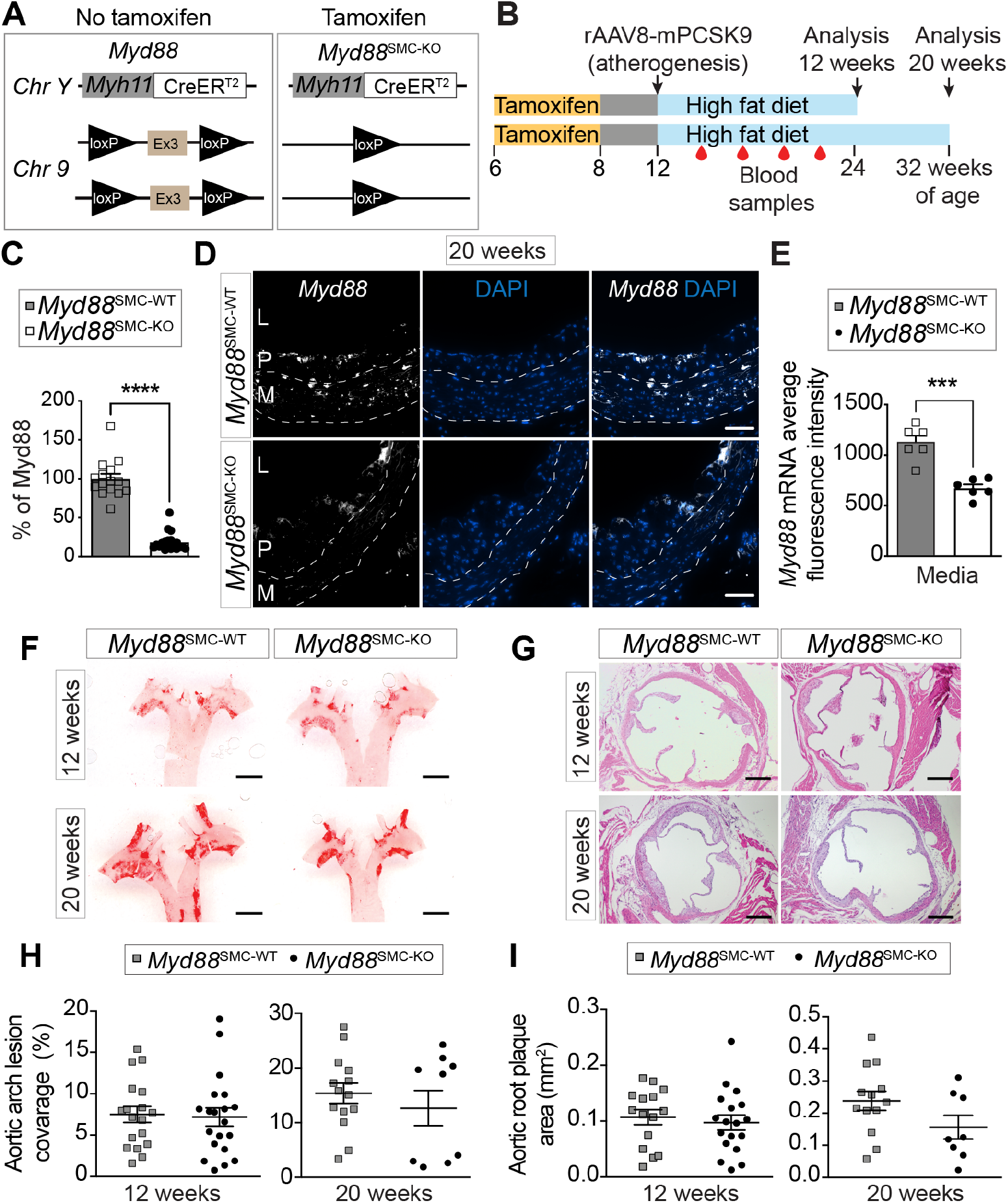
SMC-specific *Myd88* knockout does not significantly inhibit atherogenesis. **A**. Schematic illustration of the *Myh11*-CreER^T2^ x *Myd88*^flx/flx^ mouse that allows SMC-specific knockout of *Myd88* by deleting exon 3 (Ex3) after tamoxifen-induced Cre recombination. **B**. Experimental design. Tamoxifen injections (once daily for 10 days) between 6 and 8 weeks of age, followed by tail vein injection of rAAV8-mD377-mPCSK9 at 12 weeks of age and high-fat diet feeding for 12 or 20 weeks. Blood samples were collected at 2, 6, 11, and 19 weeks after rAAV8 injection. **C**. Quantitative real-time PCR for measuring the amount of remaining floxed allele after tamoxifen injections. Data shown are mean+/-SEM. Groups were compared using the Mann-Whitney test; ****P<0.0001. **D**. Representative images of RNAscope in situ hybridization for *Myd88* RNA signal (white) in aortic roots of *Myd88*^SMC-WT^ and *Myd88*^SMC-KO^ mice after 20 weeks of atherosclerosis development. Nuclei were labeled with DAPI. L (lumen), P (plaque), M (media). Scale bars, 50 μm. **E**. *MyD88* mRNA signal in the arterial media quantified as the average fluorescence intensity in media(*n*=6 mice per group). Groups were compared by unpaired t-test; ***P<0.001. **F-G**. Representative examples of *en face* examined aortic arches stained with Oil Red O and aortic root sections stained with hematoxylin-eosin in *Myd88*^SMC-WT^ and *Myd88*^SMC-KO^ mice. Scale bars, 1 mm (F) and 200 μm (G). **H**. Quantification of aortic arch lesion coverage after 12 (*n*=18-19 mice per group) and 20 weeks (*n*=9-14 mice per group) weeks of atherogenesis. No statistically significant differences detected by unpaired t-tests. **I**. Aortic root plaque area after 12 (*n*=15-18 mice per group) or 20 weeks (*n*=8-13 mice per group) of atherosclerosis development. No statistically significant differences detected by unpaired t-tests.

The mean allelic recombination frequency was measured in non-atherosclerotic aortic arterial media of the *Myd88*^SMC-KO^ mice of the 12-week experiment and found to be 83.0±1.6% (**Fig. 2C**). This translates to an estimated 64% rate of homozygous *Myd88* knockout in medial SMCs, assuming that the recombination events were randomly distributed. To confirm reduced MyD88 levels at the protein level, we tested 15 different commercial antibodies for MyD88 (as shown in **Supplemental Table 2**) but did not identify a working antibody. Cellular staining signal was obtained with several of the tested antibodies, but further testing in global *Myd88* knockout mice could not confirm specificity for any of them (data not shown). Instead, we stained aortic root sections for *Myd88* mRNA using RNAscope in situ hybridization. The analysis showed an approximately 50% reduction of Myd88 mRNA in medial ACTA2+ SMCs of *Myd88*^SMC-KO^ compared with *Myd88*^SMC-WT^ mice (**Fig. 2D-E**).

No statistically significant differences were found in final body weight in either experiment (**Supplemental Fig. 1A-B**). Nor did we detect any significant differences in plasma cholesterol levels, measured at 2, 6, 11 and 19 weeks after rAAV8-PCSK9 injection (**Supplemental Fig. 1C-D**).

### Partial loss of *Myd88* in SMCs does not reduce plaque growth or inflammation

Atherosclerosis was evaluated by *en face* examination of aortic arches and in cross-sections of the aortic root (**Fig. 2F-I**). No statistically significant difference between *Myd88*^SMC-KO^ and *Myd88*^SMC-WT^ mice was observed by either method at 12 or 20 weeks of lesion development. Since several cytokines and adhesion molecules involved in recruitment of monocytes are downstream of MyD88-dependent signaling, e.g. IL-6 and VCAM1, we also measured the abundance of LGALS3+ plaque macrophages but detected no differences between genotypes (**Fig 3A-C**). LGALS3 is expressed on a subset of SMC lineage cells in addition to macrophages, but misclassification of macrophages is modest as assessed by SMC lineage tracing,^9^ and similar findings were obtained using the more specific macrophage marker CD68 (**Supplemental Fig. 2**). Furthermore, we investigated plaque collagen content and necrotic core size and found no differences between groups (**Fig. 3D-F**).

**Figure 3.**
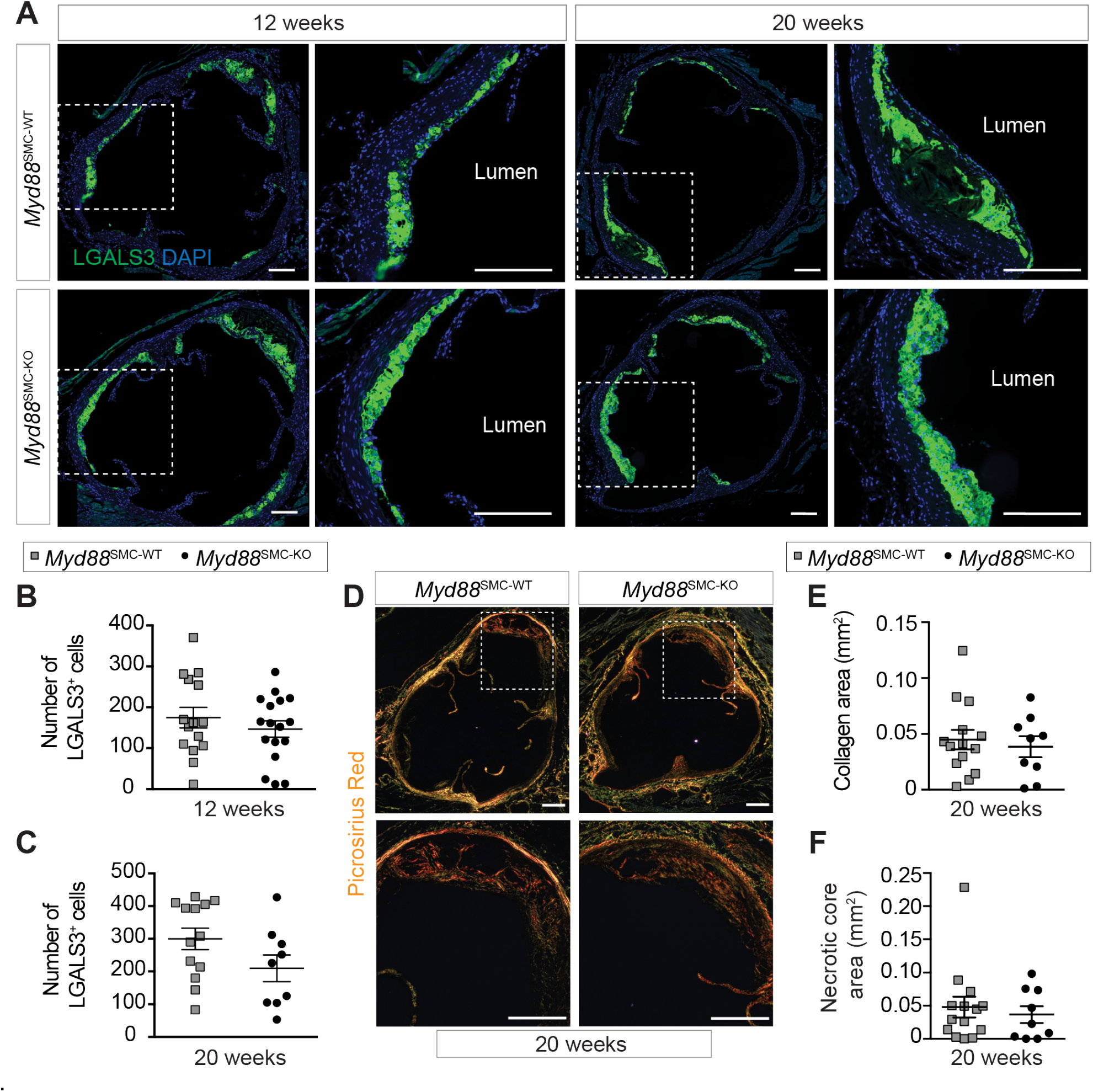
No significant changes in macrophages, collagen or necrotic core. Representative images of LGALS3-stained plaque sections from the aortic roots of *Myd88*^SMC-WT^ and *Myd88*^SMC-KO^ mice after 12 or 20 weeks of atherosclerosis development. Nuclei were labeled with DAPI. Scale bars, 200 μm. **B-C**. Number of LGALS3+ cells at (B) 12 weeks (*n*=15-17 mice per group) and (C) 20 weeks (*n*=9-13 mice per group) of atherosclerosis development. Data shown are mean+/-SEM. **D**. Representative images of picrosirius red staining of plaque sections after 20 weeks of atherosclerosis development. Scale bars, 200 μm. **E-F**. Collagen area (E) and necrotic core area (F) quantified by comparing picrosirius red-stained plaque sections from *Myd88*^SMC-WT^ and *Myd88*^SMC-KO^ mice. No statistically significant differences detected by unpaired t-tests.

### Loss of *Myd88* reduces cap SMCs

To test the main hypothesis that MyD88-dependent signaling is necessary for the recruitment of SMCs to the fibrous cap, we stained aortic root plaques for the SMC marker ACTA2, which identifies SMCs in the fibrous cap of lesions but does not stain the modified SMC-derived cells in the plaque interior.^9,25^ Changes were not detected in a combined analysis of the aortic root plaques in the three aortic sinuses after 12 weeks of atherogenesis, but after 20 weeks of atherogenesis, aortic root plaques on average had significantly fewer ACTA2+ cap SMCs in *Myd88*^SMC-KO^ compared with *Myd88*^SMC-WT^ mice (**Fig. 4A-D**). Interestingly, the loss of cap SMCs was accompanied by a statistically significant increase in ACTA2-staining cells and *Acta2* mRNA expression in the underlying media (**Fig. 4E-F**). It is well-known that many medial SMCs underlying murine plaques lose detectable ACTA2 expression. The finding that this modulation was ameliorated in *Myd88*^SMC-KO^ mice indicates that it is induced by MyD88-dependent signaling.

**Figure 4.**
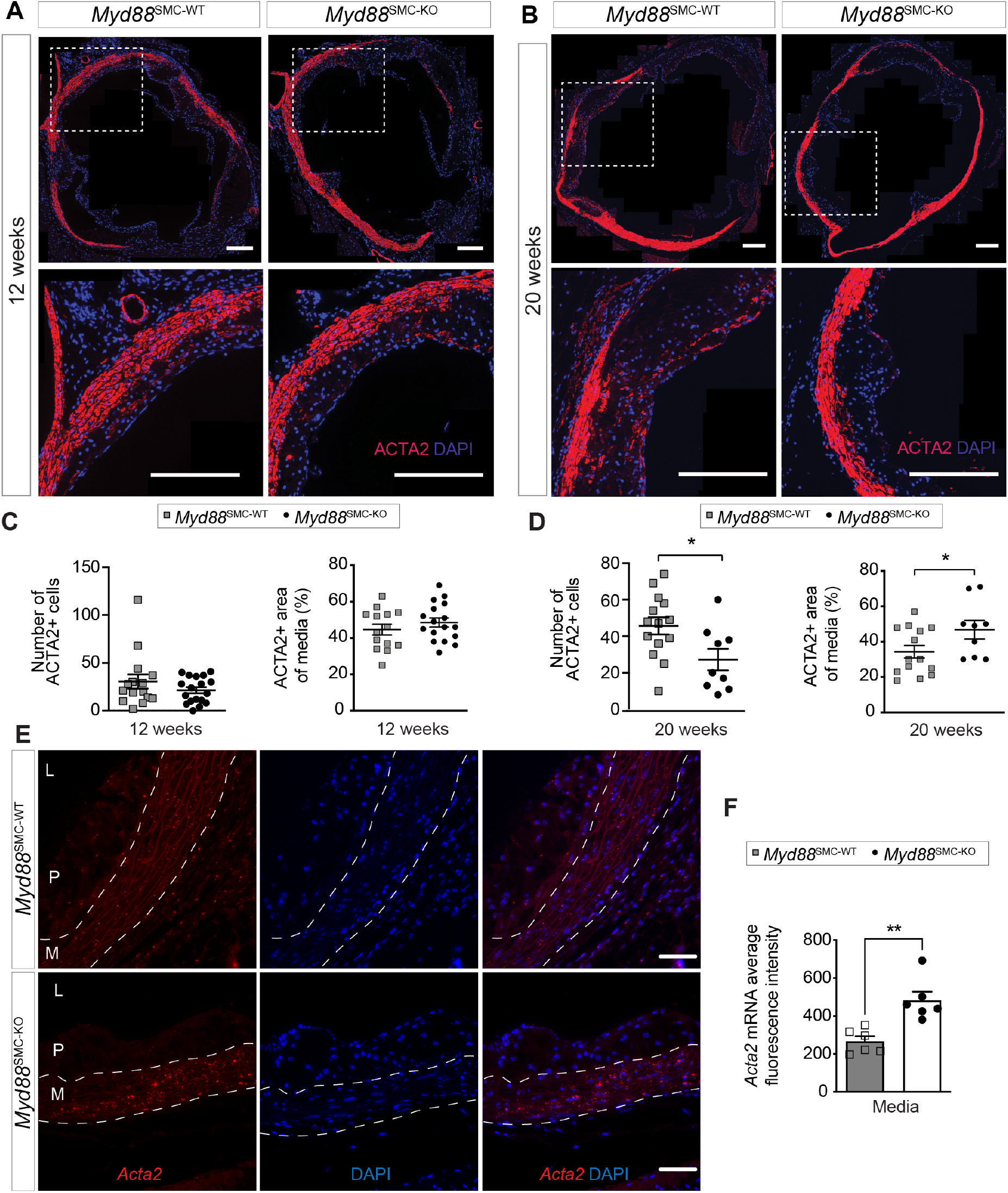
SMC-specific *Myd88* knockout inhibits cap formation and preserves medial SMC phenotype. **A-B**. Representative images of ACTA2-stained plaque sections from the aortic roots of *Myd88*^SMC-WT^ and *Myd88*^SMC-KO^ mice after 12 (A) or 20 (B) weeks of atherosclerosis development. ACTA2+ cells are mainly located in the fibrous cap. Nuclei were labeled with DAPI. Scale bars, 200 μm. **C**. Number of ACTA2+ cells in plaques and the percentage of ACTA2+ media beneath plaques after 12 (C, n=14-18 per group) and 20 (D, n=9-14 per group) weeks of atherosclerosis development. **E**. Representative images of RNAscope in situ hybridization for *Acta2* RNA signal in aortic roots of *Myd88*^SMC-WT^ and *Myd88*^SMC-KO^ mice after 20 weeks of atherosclerosis development. Each red dot corresponds to a single Acta2 RNA molecule. Nuclei were labeled with DAPI. L (lumen), P (plaque), M (media). Scale bars, 50 μm. **F**. *Acta2* RNA signal in the media quantified as the fluorescence intensity per area (*n*=6 mice per group). Groups were compared by unpaired t-tests; *P <0.05, **P <0.01.

To explore if cap SMC recruitment could be affected in the most advanced lesions already at the 12-week time point, we analyzed each aortic sinus separately. Aortic sinuses differ in atherosclerosis susceptibility, and advanced lesions develop first in the left coronary sinus. Consistently, we found that the number of ACTA2+ SMCs was already significantly reduced in left coronary sinus lesions after 12 weeks (**Supplemental Fig. 3**).

## Discussion

In the present study, we found that SMC-specific deficiency of MyD88 reduces the accumulation of cap ACTA2+ SMCs in advanced murine atherosclerotic lesions without significant changes in plaque area, necrosis, collagen content, or macrophage infiltration. This was mirrored by the preservation of contractile SMC phenotype in the underlying arterial media, indicating that MyD88-dependent signaling downstream of inflammatory cytokines or damage-associated molecular patterns causes the phenotypic modulation of medial SMCs underlying murine plaques.

The fibrous cap forms as sheets of clonal cells via PDGF and NOTCH signaling in a process that resembles - and may be conceptually related to - arterial media development.^1,3–6^ Unlike cells in the plaque interior, fibrous cap SMCs do not pass through a highly modulated LGALS3-expressing state,^26^ and they are regulated independently from SMC-derived cells in the plaque interior during plaque regression.^9^ This indicates that a distinct set of mechanisms is at play. Our finding that the process relies, at least in part, on MyD88-dependent signaling suggests that inflammatory cytokines or damage-associated molecular patterns generated in the developing atherosclerotic lesion are an important driver of recruitment and/or the expansion of fibrous cap SMCs.

This result is consistent with previous studies by Gomez et al., who investigated the role of SMC IL1R1 signaling in plaque development and fibrous cap formation in *Apoe-/-* mice.^15^ Their experiments showed that SMC-specific knockout of *Il1r1* resulted in smaller atherosclerotic lesions with fewer cap SMCs, whereas intervention with an IL-1β antibody in mice with established atherosclerosis reduced fibrous cap SMCs without altering plaque size.^15^ Differences in experimental approaches may explain why we did not detect changes in plaque area following SMC-specific *Myd88* knockout, which is expected to partially phenocopy SMC-specific *Il1r1* knockout. Atherosclerosis develops faster in the *Apoe-/-* mouse model than in mice injected with rAAV8-PCSK9, and lesions analyzed by Gomez et al. were presumably more advanced than those examined in the present study.^15^ An effect on plaque size that emerges at later stages would be consistent with the non-significant numerical decrease in plaque size we observed in *Myd88*^SMC-KO^ mice after 20 weeks of high-fat feeding.

The preservation of the contractile SMC phenotype beneath plaques in *MyD88*^SMC-KO^ mice directly recapitulated the effect of *MyD88* knockdown in cultured SMCs exposed to IL1β or LPS in the present study. Furthermore, it aligns with the well-documented pro-modulatory effect of inflammatory cytokines and NFκB signaling in SMCs *in vitro*.^7,27,28^ SMC phenotypic modulation beneath plaques is also mediated by PDGFR/LRP1 signaling, with Thymosin β4 as an upstream regulator.^29^ In vitro studies have shown that inflammation- and PDGF-induced SMC modulation occur via separate pathways,^27,28^ suggesting that the mechanism observed in the present study likely represents an independent pathway.

It is important to note that the broad phenotypic modulation that is seen beneath plaques is not necessarily sufficient for these cells to engage in the atherosclerotic process. The cells building the SMC-derived cells in murine plaque are oligoclonal, being derived from a few medial SMCs, while the media underneath the plaque typically largely stays polyclonal after labeling with stochastically recombining transgenes.^3,4^ Modulation induced by inflammatory signaling may thus be required but not sufficient for the cells to expand in atherosclerotic lesions.

### Limitations

Because this study focused on fibrous cap SMCs, for which reliable cellular markers are available, we did not perform SMC lineage tracing, which limits our ability to analyze other types of SMC-derived cells in depth. Although newer markers could potentially detect these other SMC-derived populations, the limited amount of plaque material available at the studied stage of atherosclerosis precluded such additional analyses.

## Conclusion

Overall, we found that deficiency of MyD88 preserves the contractile phenotype of medial SMCs and limits their recruitment to the fibrous cap of atherosclerotic plaques. These findings support the role of inflammatory activation of SMCs in atherosclerosis.

## Author contribution

Participated in research design: SG, JAJ, CBS, JFB

Performed experiments and data analysis: SG, JAJ, LFJ, JTS, CBS, JFB

Writing or contributing to the writing of the manuscript: SG, JAJ, CBS, JFB

Review and final approval of the manuscript: SG, JAJ, LFJ, JTS, CBS, JFB

## Acknowledgments

We thank Lisa Maria Røge and Dorte Qualmann for excellent technical assistance.

