## Supplementary Material for "MyD88-dependent signaling promotes smooth muscle cell phenotypic modulation and fibrous cap formation in murine atherosclerosis"

**Contents** **Page**

| Contents | Page |
|---|---|
| Supplemental Figure 1 | 2 |
| Supplemental Figure 2 | 3 |
| Supplemental Figure 3 | 4 |
| Supplemental Table 1 | 5 |
| Supplemental Table 2 | 6 |

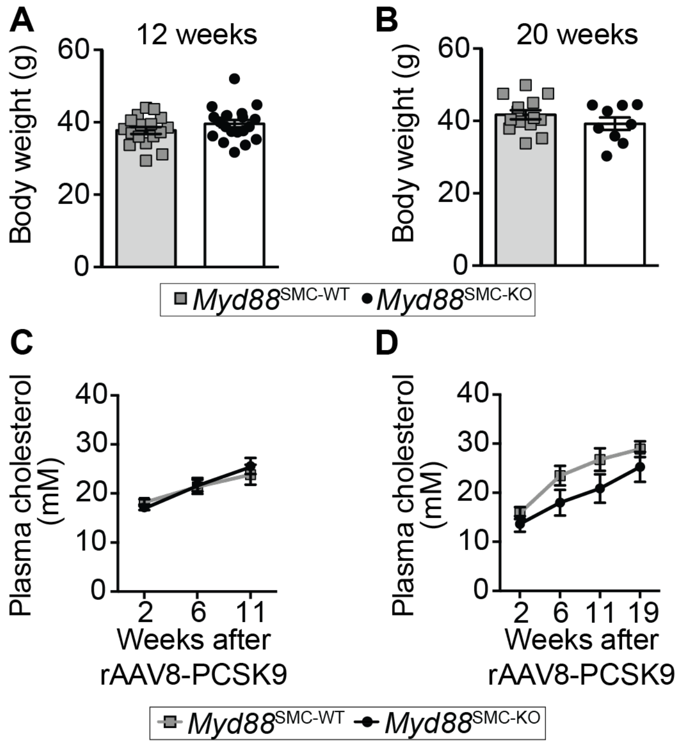


**Supplemental Figure 1. A. Body weight of *Myd88*$^{SMC-WT}$ and *Myd88*$^{SMC-KO}$ mice at (A) 12 and (B) 20 weeks of atherosclerosis development. C-D.** Plasma cholesterol levels of *Myd88*$^{SMC-WT}$ and *Myd88*$^{SMC-KO}$ mice in the 12 (C) or (D) 20 weeks experiment. Data shown are mean+/-SEM. N=9-14 mice per group. No statistically significant differences detected by unpaired t-test (A-B) and repeated-measurements ANOVA (C-D).

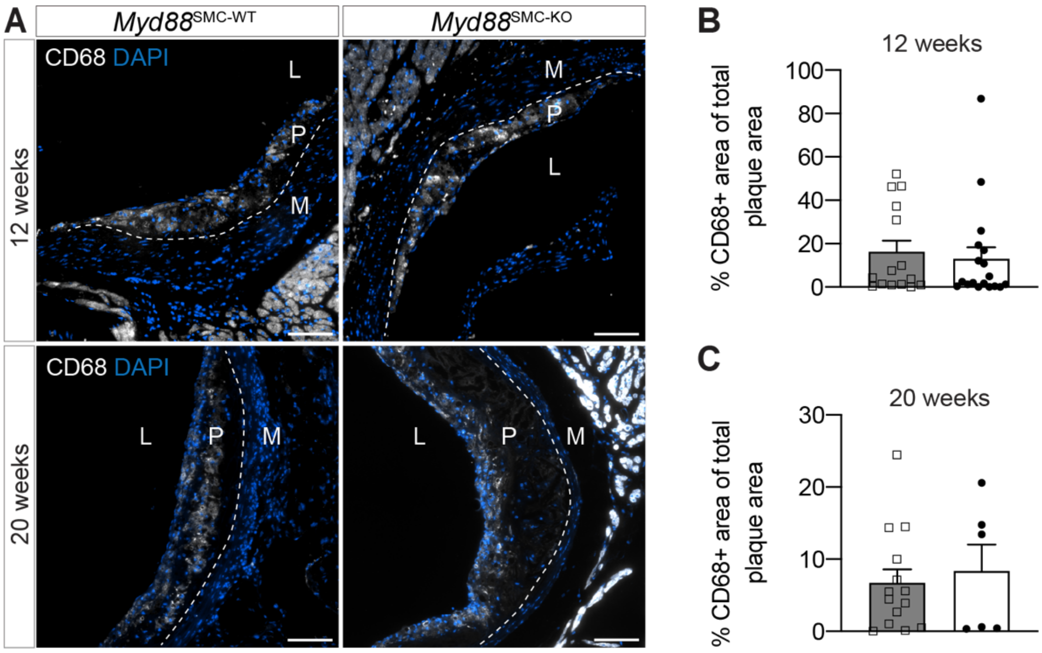


**Supplemental Figure 2. A. Representative images of CD68-stained plaque sections from the aortic roots of *Myd88*$^{SMC\text{-}WT}$ and *Myd88*$^{SMC\text{-}KO}$ mice after 12 or 20 weeks of atherosclerosis development.** Note that there is cross-reaction of the antibody to cardiomyocytes. Nuclei were labeled with DAPI. L (lumen), P (plaque), M (media). Scale bars, 100 µm. **B-C.** The percentage of CD68-positive area within the plaque area was calculated at 12 weeks (n=15-18 mice per group) and 20 weeks (n=6-14 mice per group) without detecting significant differences. Data are presented as mean ± SEM. No statistically significant differences detected by the Mann-Whitney test.

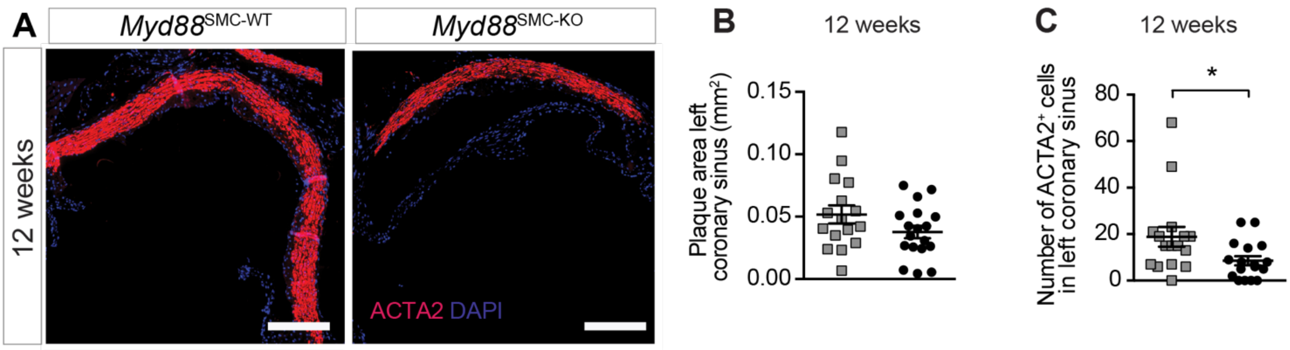


**Supplemental Figure 3. A. ACTA2 staining of the atherosclerosis-susceptible left coronary sinus of the aortic root at 12 weeks of atherosclerosis development in *Myd88*$^{SMC\text{-}WT}$ and *Myd88*$^{SMC\text{-}KO}$ mice.** Scale bars, 200 um. **B-C**. Quantification of plaque area (B) and number of ACTA2+ cells (C) in the left coronary sinus (n=16-18 mice per group). Groups were compared by unpaired t-test (B) or Mann-Whitney test (C); *P <0.05.

**Supplemental Table 1. Rat-specific primers used for qPCR analyses**

| Target gene | Primer | Primer sequence (5’-3’) |
|---|---|---|
| *Myd88* | Forward | AGAAAATGGGTCTCCCCTCA |
| | Reverse | CGAGCTCGTGCAAAGAGG |
| *Vcam1* | Forward | CCAACATGAAAGGATCGTACAG |
| | Reverse | CTCCAACAGTTCAGACGTTAGC |
| *Il6* | Forward | AGAGACTTCCAGCCAGTTGC |
| | Reverse | TGAAGTAGGGAAGGCAGTGG |
| *Myh11* | Forward | GCAGCTTCTACAGGCAAACC |
| | Reverse | GTTGATGCGAATGAACTTGC |
| *Acta2* | Forward | GCTCTGGTGTGTGACAATGG |
| | Reverse | CCCCACATAGCTGTCCTTTT |
| *Smtn* | Forward | CGGCAAGACAACAAGGAGA |
| | Reverse | ACTCCAGTCTCCCTGCCAAC |
| *Hprt1* | Forward | CTTTGCTGACCTGCTGGATT |
| | Reverse | ACTTTTATGTCCCCCGTTGA |

**Supplemental Table 2. Myd88 antibodies tested**

| Antibody: | Dilutions tested | Antigen retrival | Results |
|---|---|---|---|
| ab217141 | 1:50, 1:200, 1:500 | HIER with citrate buffer pH=6, TE-buffer pH=9 or TEG-buffer pH=9 | Identical staining in MyD88 KO tissue |
| LS-B1975 | 1:50, 1:200, 1:500 | HIER with citrate buffer pH=6, TE-buffer pH=9 or TEG-buffer pH=9 | Identical staining in MyD88 KO tissue |
| SAB2500664 | 1:50, 1:200, 1:500 | HIER with citrate buffer pH=6, TE-buffer pH=9 or TEG-buffer pH=9 | Identical staining in MyD88 KO tissue |
| ab2068 | 1:50, 1:200, 1:500 | HIER with citrate buffer pH=6, TE-buffer pH=9 or TEG-buffer pH=9 | Identical staining in MyD88 KO tissue |
| Sc-136970 | 1:50, 1:200, 1:500 | HIER with citrate buffer pH=6, TE-buffer pH=9 or TEG-buffer pH=9 | Identical staining in MyD88 KO tissue |
| Sc-74532 | 1:50, 1:100, 1:200. | HIER with citrate buffer pH=6, TE-buffer pH=9 or TEG-buffer pH=9 | No staining (TE) or identical to KO-tissue (TEG/Citrat) |
| Ab135693 | 1:10, 1:50, 1:200, 1:500, 1:1000. | HIER with citrate buffer pH=6, TE-buffer pH=9 or TEG-buffer pH=9 | Identical staining in MyD88 KO tissue |
| LS-C357983 | 1:100, 1:500, 1:1000, 1:1500 | HIER with citrate buffer pH=6, TE-buffer pH=9 or TEG-buffer pH=9 | Looks unspecific and identical in MyD88 KO tissue. |
| E-AB-32136 | 1:50, 1:200, 1:500 | HIER with citrate buffer pH=6, TE-buffer pH=9 or TEG-buffer pH=9 | Nuclear staining and identical staining in MyD88 KO tissue. |
| NBP2-67626 | 1:50, 1:200, 1:500 | HIER with citrate buffer pH=6, TE-buffer pH=9 or TEG-buffer pH=9 | Identical staining in MyD88 KO tissue |
| Ab2064 | 1:50, 1:200, 1:500 | HIER with citrate buffer pH=6, TE-buffer pH=9 or TEG-buffer pH=9 | Identical staining in MyD88 KO tissue |
| MAB 3109 | 1:50, 1:200, 1:500. | HIER with citrate buffer pH=6, TE-buffer pH=9 or TEG-buffer pH=9 | Identical staining in MyD88 KO tissue |
| TA324013 | 1:50, 1:200, 1:500. | HIER with citrate buffer pH=6, TE-buffer pH=9 or TEG-buffer pH=9 | Identical staining in MyD88 KO tissue |
| TA502117 | 1:50, 1:200, 1:500. | HIER with citrate buffer pH=6, TE-buffer pH=9 or TEG-buffer pH=9 | Identical staining in MyD88 KO tissue |
| ABIN2842312 | 1:50, 1:200, 1:500. | HIER with citrate buffer pH=6, TE-buffer pH=9 or TEG-buffer pH=9 | Identical staining in MyD88 KO tissue |
